# Low-pass filtering revisited: ramped phase randomization for cryo-EM map comparison and data augmentation

**DOI:** 10.64898/2026.10.30.763149

**Authors:** Lan Dang, Wen Jiang

## Abstract

Conventional low-pass filters attenuate or remove high-frequency Fourier amplitudes but leave the phases intact wherever the amplitude response stays above zero. As a result, fine-scale detail of the original signal can persist beyond the target resolution. A sharp cutoff removes this detail but produces ringing artifacts. To address both limitations, this work introduces ramped phase randomization, which extends a conventional low-pass filter by adding random offsets to Fourier phases that increase smoothly with spatial frequency. For cryo-EM maps, the correlation between a map at the target resolution and an ideal, noiseless map sets the levels of the phase offsets, with a correction for the finite resolution of the input map. On three published map pairs, ramped phase filtering reduced the higher resolution parent’s fine detail to a level similar to that of independently determined maps at the lower target resolution, whereas amplitude-only filtering kept the parent’s sharp features. The FSCs between the filtered maps and their parents closely resembled those of the experimental maps. The same filter also serves as a data augmentation approach for neural network-based machine learning. With ramped phase randomization, each augmented copy keeps the power spectrum and low-frequency content of the original example but has a different high-frequency pattern. For simulated CryoBench particles of 16 ribosome assembly states, phase augmentation raised mean test accuracy in all 20 combinations of training-set size and latent dimension, by 20.8 percentage points on average (range from 3.3 points to 44.2 points). On two general deep learning benchmarks, mean test accuracy rose from 92.15% to 96.99% with 60 labelled UCI Synthetic Control traces and from 53.44% to 74.27% with 500 labelled noisy MNIST images. The ramped phase randomization is implemented in an open-source Python package, with a companion browser application for interactive visualization of cryo-EM maps and filtering effects.

**Synopsis:** Combining amplitude filtering with ramped phase randomization reduces the fine-scale phase information after low-pass filtering, with applications to cryo-EM map comparison at a common resolution and to data augmentation for more robust learning.

## 1. Introduction

Every measurement has a resolution limit, beyond which high-frequency contributions are dominated by noise or cannot be determined reliably. Low-pass filtering in Fourier space is commonly used to suppress these contributions beyond a chosen target resolution.

Conventional low-pass filters take two main forms (Gonzalez & Woods, 2018). The first form is band limitation, which sets Fourier coefficients beyond a cutoff to zero, removing both amplitude and phase. This sharp cutoff in Fourier space, however, is equivalent to convolving the signal with an oscillatory, sinc-like kernel in real space, which produces ringing artifacts around features. The second form is smooth amplitude attenuation, which reduces ringing by lowering coefficient magnitudes progressively with frequency. Wherever the amplitude response stays above zero, this filter leaves the phases intact. However, phases encode much of the spatial or temporal organization of a signal (Oppenheim & Lim, 1981). Figure 1A illustrates the role of phases in a 2D signal: when two images exchange Fourier amplitudes and phases, each hybrid resembles the image that supplied its phases. Therefore, amplitude filtering only weakens but does not remove fine-scale detail beyond the target resolution.

**Figure 1.**
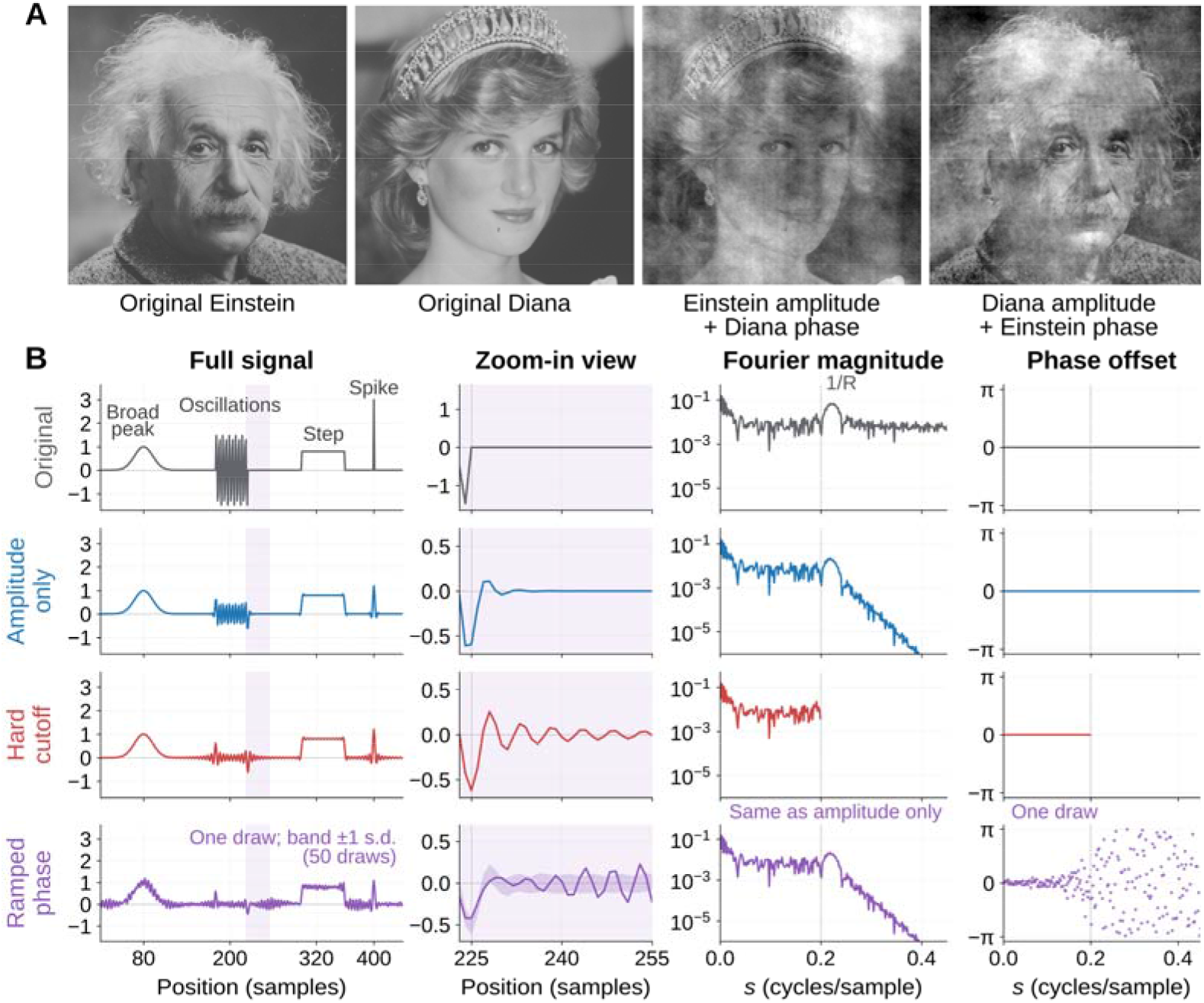
Phase exchange and filtering. (A) Phase-swapped portraits largely retain the facial features of the phase donor. (B) Amplitude-only, hard-cutoff, and ramped-phase filtering to *R* = 5 samples, with zoomed views of the dip and ringing. differs between realizations (Figure 1B).

To limit the phase information that amplitude filtering leaves beyond the target resolution, we develop a general low-pass operator that combines amplitude attenuation with random phase offsets. The spread of the offsets ramps up smoothly with frequency rather than switching on at a cutoff, so we call this approach ramped phase randomization. This ramped operator leaves low-frequency components largely unchanged, whereas higher-frequency components progressively lose correlation with the original signal, which we call the parent. We demonstrate two areas of applications of the operator: comparing cryo-EM maps at a common resolution, and augmenting training sets for machine learning.

In cryo-EM, maps of the same macromolecular complex may differ in resolution because the maps come from different datasets, conformational states or reconstruction methods. Comparing such maps requires separating structural differences from differences introduced by resolution and image processing. Common workflows align the maps, filter both to a common resolution and match the Fourier amplitudes before calculating a difference map (Joseph *et al*., 2020; Warshamanage *et al*., 2022). However, amplitude filtering alone keeps the phases of the higher-resolution parent map beyond the target resolution. The filtered map therefore retains fine detail that the lower-resolution map does not reliably resolve. This residual detail can produce features in the difference map that mimic structural changes. Limiting this detail requires changing the phases, not only the amplitudes, beyond the target resolution.

Phase randomization beyond a fixed frequency was introduced in cryo-EM as a test for overfitting (Chen *et al*., 2013; see Sorzano *et al*., 2022 for a critical discussion). We instead use phase randomization as a filter and schedule the randomization by spatial frequency. At each frequency, we set the spread of the phase offsets according to the correlation between a map at the target resolution and an ideal, noiseless map (Rosenthal & Henderson, 2003). Because the parent in practice is itself imperfect, the offsets add only the remaining loss of correlation. The filtered map is then expected to agree with the ideal map as closely as one determined at the target resolution. We use the Fourier shell correlation (FSC; Harauz & van Heel, 1986) to measure how closely the filtered map still agrees with the parent. We first compare this FSC with that of experimental maps, using three published pairs of high- and low-resolution maps of the same complex. We then examine ribosome difference maps, using a comparison within the same conformational state as a control for a comparison between states.

The same ramped phase operator also addresses a common problem in machine learning. Neural networks with many trainable parameters can memorize details and noise specific to individual training examples, and so generalize poorly to unseen data, a problem known as overfitting (Goodfellow *et al*., 2016; Arpit *et al*., 2017). Data augmentation can reduce overfitting by varying features that are irrelevant to the task while preserving those needed for prediction (Shorten & Khoshgoftaar, 2019; Iwana & Uchida, 2021).

Several methods perform data augmentation in the Fourier domain. For example, some image augmentation schemes swap the amplitude spectra of different images while retaining the original phases (Chen *et al*., 2021; Xu *et al*., 2021). These augmentations train a model to rely on image structure rather than image style, which makes the model more robust to domain shifts and image corruptions. Alternatively, some data augmentation schemes perturb phases equally at all frequencies while keeping the power spectrum. Such phase randomization generates surrogate time series (Theiler *et al*., 1992; Lancaster *et al*., 2018), which can enlarge small or imbalanced training sets of physiological recordings such as electroencephalography signals (Schwabedal *et al*., 2019; Rommel *et al*., 2022). However, for tasks that depend on waveform shape, these approaches involve a trade-off. Preserving all phases can retain the noise pattern of each example, whereas perturbing phases across the entire spectrum can disrupt informative low-frequency structure.

We address this trade-off with the same ramped phase randomization, applied without changing the amplitudes. Instead of following a target resolution, the offsets start at a chosen frequency and grow toward the high end of the spectrum, so the phases below that frequency are kept. Each augmented copy therefore preserves the original power spectrum and low-frequency content but carries a new high-frequency pattern. We hypothesize that, when the information needed for classification lies mainly at low frequencies, this variation pushes a model to learn the shared low-frequency signal rather than example-specific noise, thereby improving generalization to new examples.

Cryo-EM particle images provide a natural first test, because the images are noisy and labelled examples are often scarce. We use simulated particle images of 16 ribosome assembly states, viewed from random directions, from the Ribosembly dataset in CryoBench (Jeon *et al*., 2024) and train a classifier to identify the assembly state of each particle. To test whether the augmentation also applies beyond cryo-EM, we use two general machine-learning benchmarks in which the class information lies mainly at low frequencies: the UCI Synthetic Control Chart time series (Alcock, 1999), and MNIST digits (LeCun *et al*., 1998). In all three cases, we systematically vary the training-set size and the latent dimension of the network, and measure how much augmentation reduces overfitting in each setting.

## 2. Theory: Low-pass filtering of amplitudes and phases

### 2.1 Limiting retained phase information

Let *F*_*H*_(**k**) denote the discrete Fourier transform of a signal *H* sampled on a *d*-dimensional grid, such as a time series (*d* = 1), an image (*d* = 2) or a cryo-EM density map (*d* = 3), with frequency magnitude *s* = |**k**|. Each Fourier coefficient can be written in the amplitude–phase form as 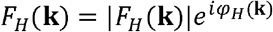, where the amplitude |*F*_*H*_ (**k**)| sets how strongly the corresponding sinusoid contributes to the signal and the phase *φ*_*H*_(**k**) sets the position of that sinusoid. A low-pass filter limits a signal to a chosen resolution *R* by keeping coefficients well below the frequency 1/*R* and attenuating or removing those above 1/*R*. Applied to the parent signal *H*, a conventional low-pass filter multiplies each coefficient by a real, non-negative amplitude response, 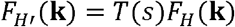. The filter changes the amplitudes but leaves the phase *φ*_*H*_ (**k**) intact wherever *T*(*s*) > 0. The retained phases can be observed clearly in three-dimensional cryo-EM maps, where the agreement between two maps *X* and *Y* is measured by the FSC over the coefficients in a thin spherical shell *S*_*s*_ of radius *s*:

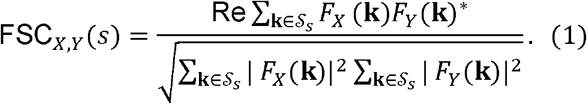

For an amplitude-only filter, the FSC with the parent stays at unity however strongly the amplitudes are attenuated, because a positive amplitude response constant within a shell cancels between numerator and denominator. The same normalization can make strongly low-pass-filtered maps appear highly correlated, a known pitfall of FSC analysis (Van Heel & Schatz, 2005). A unit FSC means that the filtered map keeps every parent phase beyond the target resolution, leaving fine structure in place but weaker. Rescaling the amplitudes, as B-factor sharpening does (Rosenthal & Henderson, 2003; Fernández *et al*., 2008), could in principle restore that structure.

A low-pass filter that limits the information retained beyond the target resolution should therefore modify the phases as well as the amplitudes. Such a general operator can be written as:

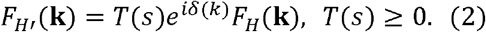

Where the amplitude response *T*(*s*) scales the amplitude of each coefficient, as in a conventional low-pass filter, and the random phase offset *δ*(**k**) shifts the phase from *φ*_*H*_(**k**) to *φ*_*H*_(**k**) + *δ*(**k**), disrupting the relative positions of features that the filtered signal inherits from the parent. For offsets drawn from a distribution that is symmetric about zero and depends only on the frequency magnitude *s*, the first circular moment, *c*(*s*), which we call the coherence, is real and measures how closely the filtered phases follow the parent:

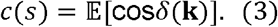

With no phase perturbation, *c* = 1; for the wrapped-normal offsets used here (Section 2.2), broader offsets reduce *c* toward 0. The phase perturbation appears directly in the FSC of equation (1): for a shell *S*_*s*_ in which *T*(*s*) is positive and constant, and in which all offsets follow the same distribution,

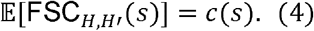

A gradual increase of the offsets with frequency produces a smooth decline in the FSC. Figure 1B compares the filters on a one-dimensional signal. Amplitude-only filtering leaves an attenuated copy of a high-frequency oscillation (Figure 1B, full signal). A hard cutoff filter removes the oscillation but adds periodic ringing where the original signal is zero, as the zoom-in view shows after the oscillation ends at sample 225. Ramped phase filtering keeps the same Fourier magnitudes as amplitude-only filtering (Figure 1B, Fourier magnitude) but randomizes the phases above the target frequency (Figure 1B, phase offset). The oscillation is then replaced by a noise pattern that spreads along the signal instead of ringing at the edge, and that

### 2.2 A coherence envelope that mimics correlation with an ideal signal

The ramped phase randomization is specified by choosing the coherence profile c(s) in equation (3). We set the profile to the correlation that a signal at resolution *R* would share with an ideal, noiseless signal. By equation (4), the filtered signal then correlates with an ideal parent as strongly as a signal at resolution *R* would. This correlation is near one at low frequency and falls through the target band. We adopt the convention that a signal at resolution R has a correlation of 0.5 with the ideal signal at frequency 1/R, which for cryo-EM maps corresponds to the half-map FSC = 0.143 criterion (Rosenthal & Henderson, 2003). For an ideal parent, expressing the correlation through the spectral signal-to-noise ratio (SSNR) and modelling the SSNR with a Gaussian falloff gives

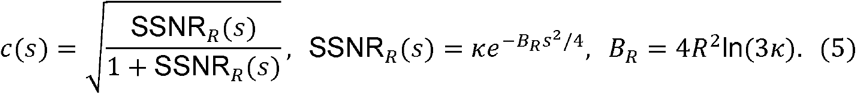

Here *R* is a length (Å) for density maps, samples for time series), *s* has the reciprocal unit and *B*_*R*_ the squared unit; *κ* is dimensionless. A large *κ*is chosen so that the profile stays near unity at low frequency. The constant *B*_*R*_ fixes the crossing so that c(1/*R*) = 0.5. Each phase offset is drawn at random from a normal distribution centred on zero, so that shifts in either direction are equally likely. Because adding 2 Π to a phase leaves a Fourier coefficient unchanged, offsets larger than Π simply wrap around the circle. The standard deviation *σ*(s) of the underlying normal distribution sets *c*:

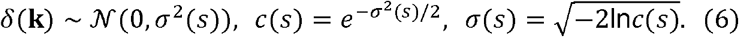

The third relation gives the width needed for a chosen coherence (derivation in Supplementary Section S1). With *c* from equation (5), the offsets are small at low frequency and grow progressively through the target-resolution band. The amplitude response is initially set to *T* = *c* and may later be rescaled, for example by radial amplitude matching (Section 5.2), without changing the phase offsets. The 0^th^ Fourier coefficient (DC), which sets the mean of the signal, is excluded from filtering. The ramping envelope is a model, not a noise fit: a measured coherence profile can replace the envelope where available.

This ramped construction makes the expected filtered signal fall off smoothly through 1/*R*. Binary phase randomization, such as high-resolution noise substitution (Chen *et al*., 2013), instead keeps *c* at one up to 1/*R*, where a signal at resolution *R* correlates with the ideal signal at only 0.5. With the amplitude response of equation (5), the expected filtered signal drops abruptly from half amplitude to zero at 1/*R*. This step produces ringing that survives averaging over random draws.

### 2.3 Accounting for phase uncertainty already present in the parent

The parent already differs from the ideal signal, so the resolution of the filtered signal depends on the parent. Let *a*_*H*_(s) denote the parent envelope, the correlation of the parent with the ideal signal, and let *a*_*T*_(s) denote the target envelope. Phase randomization multiplies this correlation by *c*(*s*), so the filter only needs to add the ratio of the target envelope to the parent envelope, capped at one:

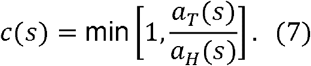

For an ideal parent, the parent envelope equals one and equation (7) reduces to equation (5). In practice, both envelopes come from equation (5), evaluated at the parent resolution *R*_*H*_ and at the target resolution *R*_*T*_. With the same *κ*, the target envelope lies below the parent envelope at all frequencies whenever *R*_*T*_≥ *R*_*H*_. Before radial matching, the amplitude response is set to the same ratio, *T* = *c*. Frequencies where the parent envelope is zero are discarded. At the target frequency, the filtered signal correlates with the ideal signal at 0.5, as defined in Section 2.2. Equation (7) describes a single filtered map, the form in which the filter is used; averaging realizations would restore agreement with the parent (Supplementary Figure S1D, E). By contrast, two independently determined maps and of the same structure, with uncorrelated errors, have an expected FSC equal to the product of their envelopes rather than the ratio (Supplementary Section S1):

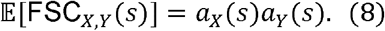

Half maps are two such maps of equal quality, so their FSC *ρ*,_1/2_, equals 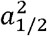. Their average, the full map, has half the error power and envelope 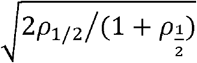, so a half-map FSC of 0.143 corresponds to a correlation of 0.5 between the full map and the ideal map, the criterion of Section 2.2 (Supplementary Figure S1A, B). This generalizes the relation between half-map FSC and map quality (Rosenthal & Henderson, 2003; Agarwal *et al*., 2024) to maps of different quality. With envelopes from equation (5), equation (8) predicts the FSC between two maps from their nominal resolutions (Supplementary Figure S1C). At the target frequency, the expected FSC with the parent is thus slightly below 0.5 for an independent map at the target resolution and slightly above 0.5 for the filtered map. For experimental data such as cryo-EM maps, nominal resolutions only approximate the true frequency-dependent phase reliability.

## 3. Results

Sections 3.1 and 3.2 test the ramped phase randomization operator on three-dimensional cryo-EM density maps, where the FSC (equation 1) measures the agreement retained with the parent after filtering and independently determined maps of the same complex provide experimental references. Section 3.3 applies the same operator as a training data augmentation for neural network-based machine learning, first to simulated cryo-EM particle images and then to one- and two-dimensional benchmark signals.

### 3.1 Ramped phase filtering removes parent detail beyond the target resolution

To test the filter on experimental data, each of three higher-resolution maps, of T20S proteasome (Campbell et al., 2015; Huber et al., 2022), the *Saccharomyces cerevisiae* 80S ribosome in situ (Rangan *et al*., 2024; Weber *et al*., 2026) and apoferritin (Kayama et al., 2021), was filtered to the resolution of an independently determined lower-resolution map of the same complex (Figure 2), which provides an experimental reference at the target resolution. Each pair was registered to a common grid (Section 5.1), and equation (7) corrected each parent for nominal resolution before radial amplitude matching to the lower-resolution map (Section 5.2). As a control, each parent was also filtered with the same amplitude response but without phase offsets (amplitude-only filtering). The target resolution of each pair was the half-map FSC = 0.143 resolution of the lower-resolution map, as calculated by the EMDB without masking. At that frequency, the full map is expected to have a correlation of 0.5 with the ideal structure (Rosenthal & Henderson, 2003; Section 2.3), the criterion that defines R in Section 2.2.

**Figure 2.**
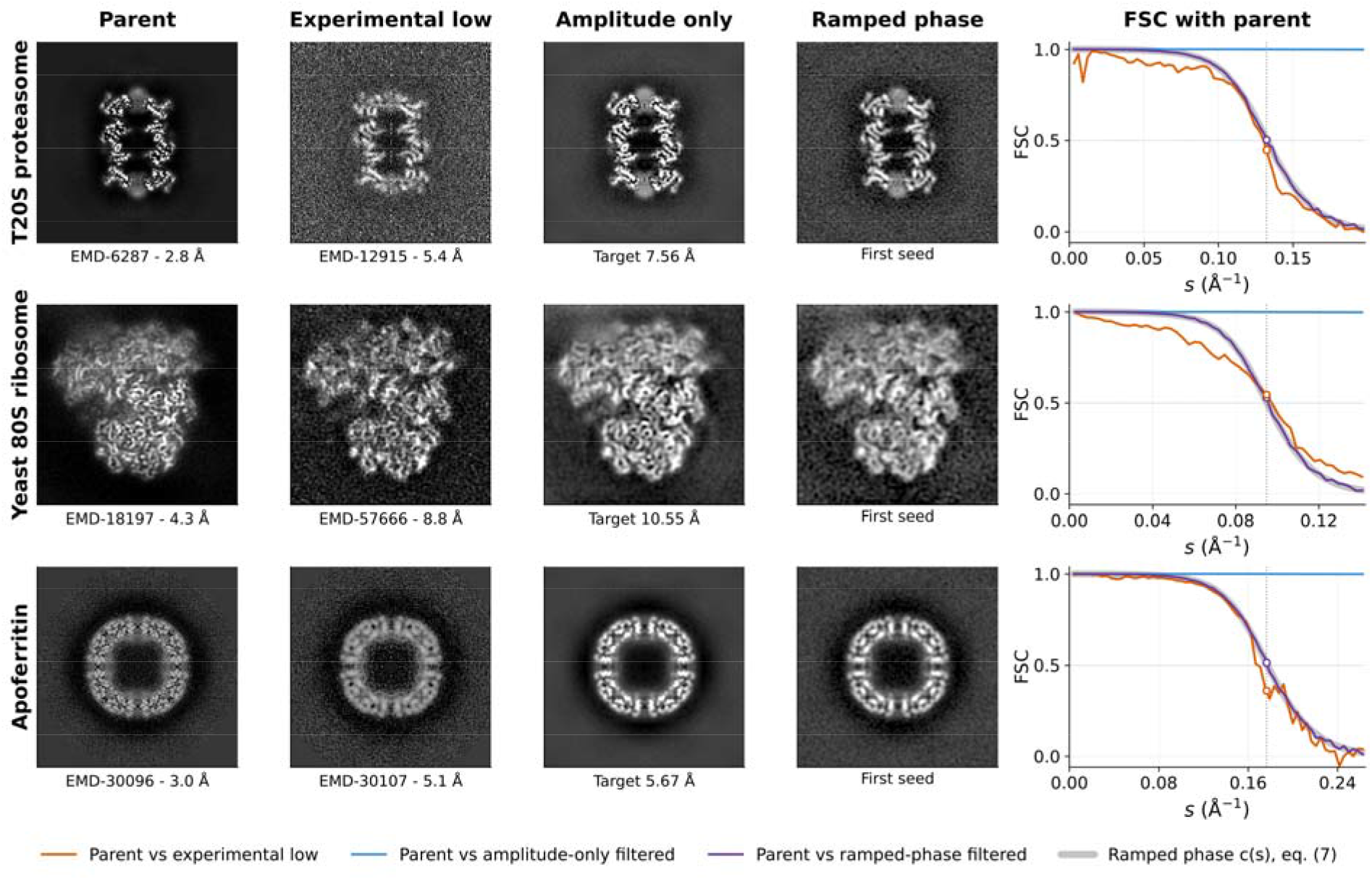
Published map pairs after amplitude-only and ramped-phase filtering. Rows show T20S proteasome, the 80S ribosome in situ and apoferritin. Columns show central sections of the registered parent, the experimental lower-resolution map, the amplitude-only map and the ramped-phase map (first of 50 seeds), followed by FSCs with the parent. Targets (vertical lines) are the EMDB-calculated unmasked half-map resolutions of the lower-resolution maps. Both filters use the same parent-corrected amplitude response; ramped-phase filtering adds wrapped-normal phase offsets with coherence from equation (7). Gray bands: the ramp c(s) of equation (7), the expected FSC of the ramped-phase map; circles mark the FSCs at the target. All FSCs are unmasked.

Both filters kept the low-resolution structure of the parent, but only phase filtering suppressed the finer detail, increasingly through the target-resolution band. Visually, the ramped-phase maps resembled the lower-resolution experimental maps in their level of noise and fine detail, whereas the amplitude-only maps retained more sharp features from the parent (Figure 2). The FSC with the parent quantifies the difference (Figure 2, final column). The amplitude-only FSC stayed at 1 at all frequencies, even where amplitudes were strongly reduced (Section 2.1), whereas ramped phase filtering lowered the FSC smoothly. At the target frequency, the ramped-phase maps followed equation (7) (gray bands in Figure 2): over 50 seeds, the mean FSC was within 0.002 of the expected value of about 0.51 for each pair, and individual seeds ranged from 0.49 to 0.53. For comparison, equation (8) predicts an FSC of 0.49 for an independent map at the target resolution, close to the 0.51 of the filtered maps. The prediction is nearly the same for all three pairs because each parent is two to three times higher resolution than its target and so agrees almost perfectly with the ideal map at the target frequency. The experimental maps scattered around this value, from 0.36 to 0.55, because independent maps also differ in ways their nominal resolutions do not capture, such as microscope optics, sample preparation and image processing (Section 2.3). The filtered and experimental FSC curves had substantially similar shape, crossing 0.5 within 0.3 Å of each other for all three pairs.

### 3.2 Cross-resolution difference maps localize ribosome inter-subunit rotation

Two cryo-EM density maps of the same complex determined at different resolutions are difficult to compare directly. A rigorous comparison should involve three steps: registration, which resamples both maps onto a common grid, and aligning scale, orientation and position (Section 5.1); radial amplitude scaling, which matches the Fourier shell-mean amplitudes of the two maps; and phase randomization of the high-resolution map to the level expected at the lower resolution (Section 5.2).

We tested this approach with a 3.0 Å *Escherichia coli* 70S ribosome map (Carbone et al., 2021) as the parent and two 13.2 Å pre-translocation maps from an earlier study, one without relative rotation of the large and small subunits (non-rotated) and one rotated (Fu et al., 2011). The non-rotated map shares the conformational state of the parent and serves as a control. The parent was registered separately to each of the two target maps; the two transforms agreed to within 0.23% in scale, 0.64° in orientation and 0.40 Å in translation, so both comparisons used nearly the same registration.

After filtering to 13.2 Å, the parent agreed less with the rotated map than with the non-rotated control: at the shell nearest the target (0.0747 Å^−1^), the FSC was 0.197 ± 0.005 against the rotated map and 0.350 ± 0.009 against the control (mean ± SD over eight seeds), close to the predicted values of 0.197 and 0.347, obtained by multiplying the unfiltered FSCs, 0.368 and 0.649, by = 0.534 (Supplementary Section S2). The difference maps show where the disagreement lies. The control difference was weaker and more diffuse (Figure 3D, E), whereas the rotated difference contained a strong, localized signal near the subunit interface (Figure 3H, I), consistent with inter-subunit rotation. Inside the molecular region, the residual variance of the rotated comparison was about 2.5-fold that of the control (Section 5.3). Individual phase-randomization seeds gave nearly identical results (2.45–2.48-fold across eight seeds), so Figure 3 shows a single realization.

**Figure 3.**
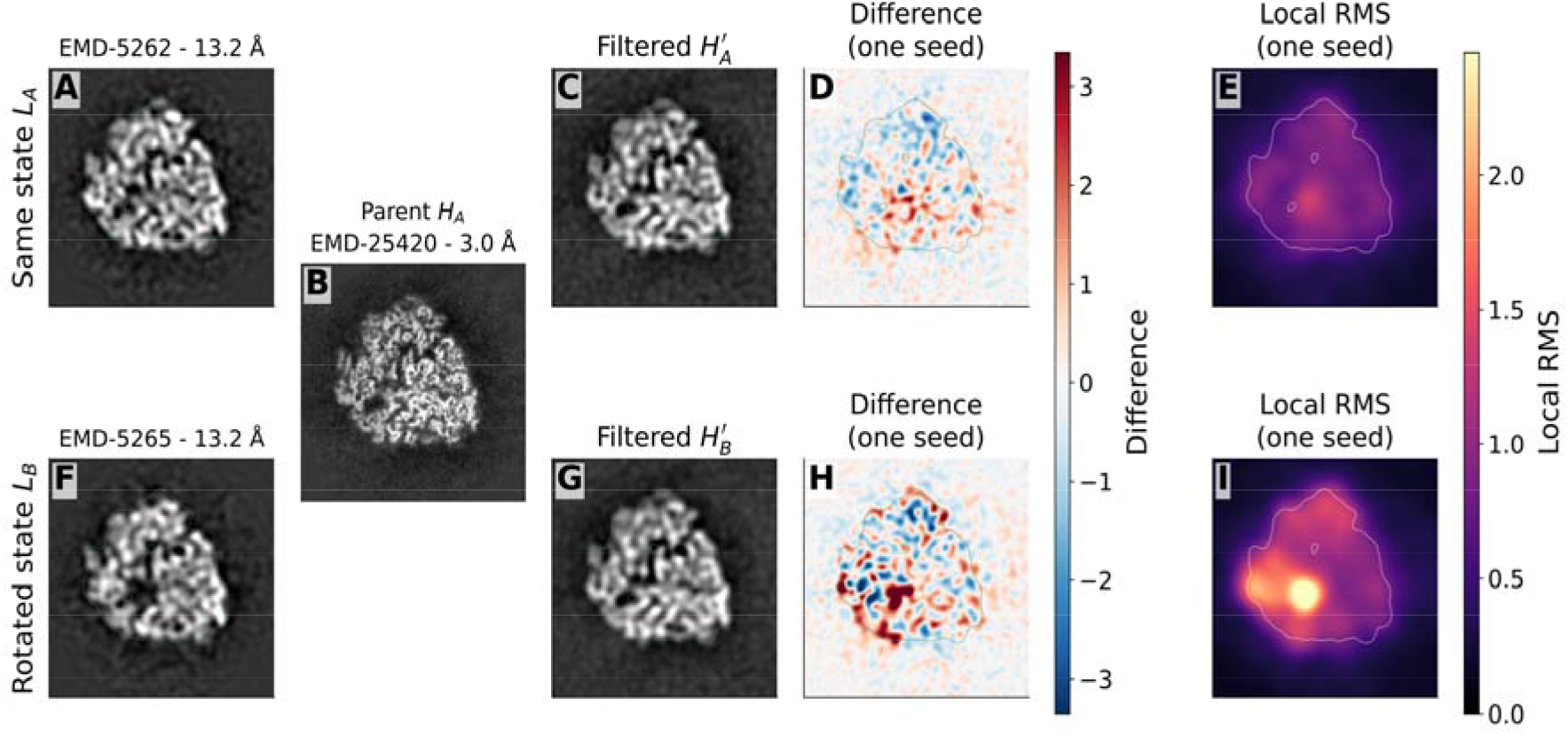
Ribosome difference maps for the non-rotated control (top) and the rotated state (bottom). (A, F) Experimental 13.2 Å maps *L*_A_ and *L*_B_. (B) Parent *H*_A_ registered to *L*_A_, on the same plane as A and C. (C, G) Parent filtered for each comparison (*H*′_A_, *H*′_B_; one phase-randomization seed). (D, H) Signed differences between the filtered and experimental maps. (E, I) Local rms of the differences (Gaussian kernel, FWHM 31 Å), on a shared scale.

The control difference was not zero. The parent and control come from different acquisitions, contain different tRNAs and were processed differently, and the control specimen carries a P-loop mutation. The comparison thus highlights where the rotated map departs from the parent more than an independent map of the same state does.

### 3.3 Phase randomization as data augmentation for noise-dominated data

The second area of applications uses a ramped phase randomization operator as a training augmentation for small, noisy datasets. Such datasets are common in experimental science, for example when cryo-EM particle pickers are trained on a few manually picked particles (Bepler *et al*., 2019).

We first tested whether phase augmentation improves generalization of supervised learning on noisy, simulated cryo-EM particle images. Particle images were from the Ribosembly dataset in CryoBench (Jeon *et al*., 2024), which contains simulated projections from 16 maps, each representing one ribosome assembly state (Qin *et al*., 2023; Figure 4, top rows). We used the deposited images, with their random viewing directions, CTF modulation and nominal signal-to-noise ratio of 0.01. We used the poses supplied with CryoBench instead of estimating them. Because these are the poses used to simulate the images, the setup mimics an ideal consensus refinement. Each image was shifted to the box centre, phase-flipped with its supplied CTF and backprojected along its viewing direction onto a 2 Å 3D grid, fine enough to keep every frequency of the 3 Å images (Section 5.4). This would generate a single-view backprojection 3D grid that coherently encodes the projection image and the corresponding pose. A classifier assigned each of these grids to one of the 16 states through a low-dimensional latent layer (Supplementary Table S1). Training sets ranged from 16 to 256 particles per state and latent dimensions from 2 to 16. Sweeping both factors shows how the benefit depends on the amount of labelled data and on latent capacity.

**Figure 4.**
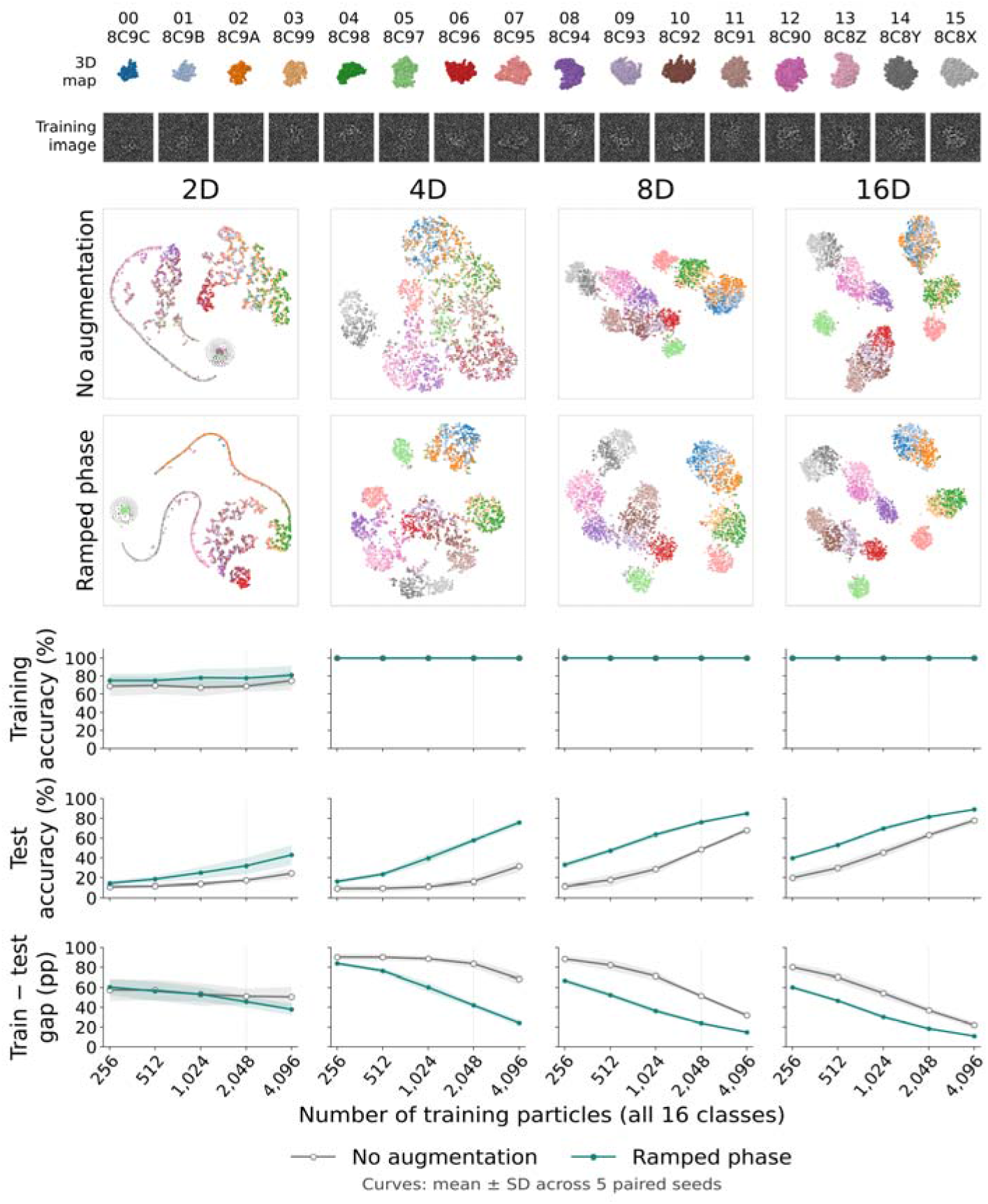
Ramped phase augmentation of CryoBench particle images. Top: the 16 Ribosembly maps, labelled with the source PDB IDs (Qin *et al*., 2023), and one training image per state (112 × 112 crop). Middle: t-SNE projections of the latent features of 4,096 test particles, coloured by state, from one paired seed trained on 128 particles per state (dotted lines), with separate latent dimensions in each column. Bottom: training accuracy, test accuracy and train–test gap against training-set size for 2-, 4-, 8- and 16-dimensional latent layers, without (grey) and with (teal) augmentation, as mean ± SD over five paired seeds.

During training, each CTF-corrected image was augmented with probability 0.5 before backprojection. Each augmentation drew a random start frequency between 0.036 and 0.044 cycles per pixel (83–68 Å), with the end frequency fixed at 0.20 cycles per pixel (15 Å). We trained five paired seeds; within each seed, the two conditions, with or without phase randomization augmentation, shared the same training particles, initial weights and minibatch order.

Phase augmentation raised mean test accuracy in all 20 settings and in 99 of 100 individual seeds (Figure 4). It also improved the grouping of test particles by state in all 20 settings, measured by the adjusted Rand index (ARI; Hubert & Arabie, 1985) of k-means clusters (Supplementary Section S3). With an eight-dimensional latent layer and 128 particles per state, test accuracy rose from 48.5 ± 1.8% to 76.1 ± 1.7% (mean ± SD over five seeds; paired gain 27.6 percentage points, 95% interval 25.9–29.2), and the ARI rose from 0.347 to 0.565. Across the sweep, mean gains ranged from 3.3 to 44.2 percentage points (average 20.8), and augmented training beat unaugmented training on twice as many particles in all comparisons. From four latent dimensions upward, both conditions fitted the training set well, and augmentation reduced the train–test gap in every setting; with two latent dimensions, neither condition fitted the training set well (67–81% training accuracy). In t-SNE projections of the latents (van der Maaten & Hinton, 2008), test particles of different states overlapped more without augmentation and formed clearer groups with it, most visibly at 4 and 8 dimensions (Figure 4, middle rows).

To test whether the benefit extends beyond cryo-EM, we also compared phase augmentation with no augmentation on two general machine-learning benchmarks, using 25 paired seeds for each (Supplementary Section S3). The UCI Synthetic Control Chart dataset contains short time series in six classes defined by the presence and type of trend, shift or cycle (Alcock, 1999). Most class-defining power lies at low frequencies, and most high-frequency content consists of fluctuations specific to each trace (Figure 5A). A classifier with an eight-dimensional latent layer was trained on 60 labelled traces, 10 per class, and tested on 300 unseen traces. Phase augmentation raised mean test accuracy from 92.15% to 96.99% (mean paired gain 4.84 percentage points, 95% interval 3.93–5.75), with higher test accuracy in all 25 seeds, while training accuracy stayed at 100% in both conditions (Figure 5B). The gain therefore came from better generalization rather than better fitting. Without augmentation, most errors came from three pairs of similar classes: normal and cyclic traces, increasing trends and upward shifts, and decreasing trends and downward shifts. Pooled over seeds, phase augmentation reduced the errors within each pair by 44–63%, most strongly between normal and cyclic traces, and separated the same pairs in the hidden features of an illustrative model (Figure 5C, D).

**Figure 5.**
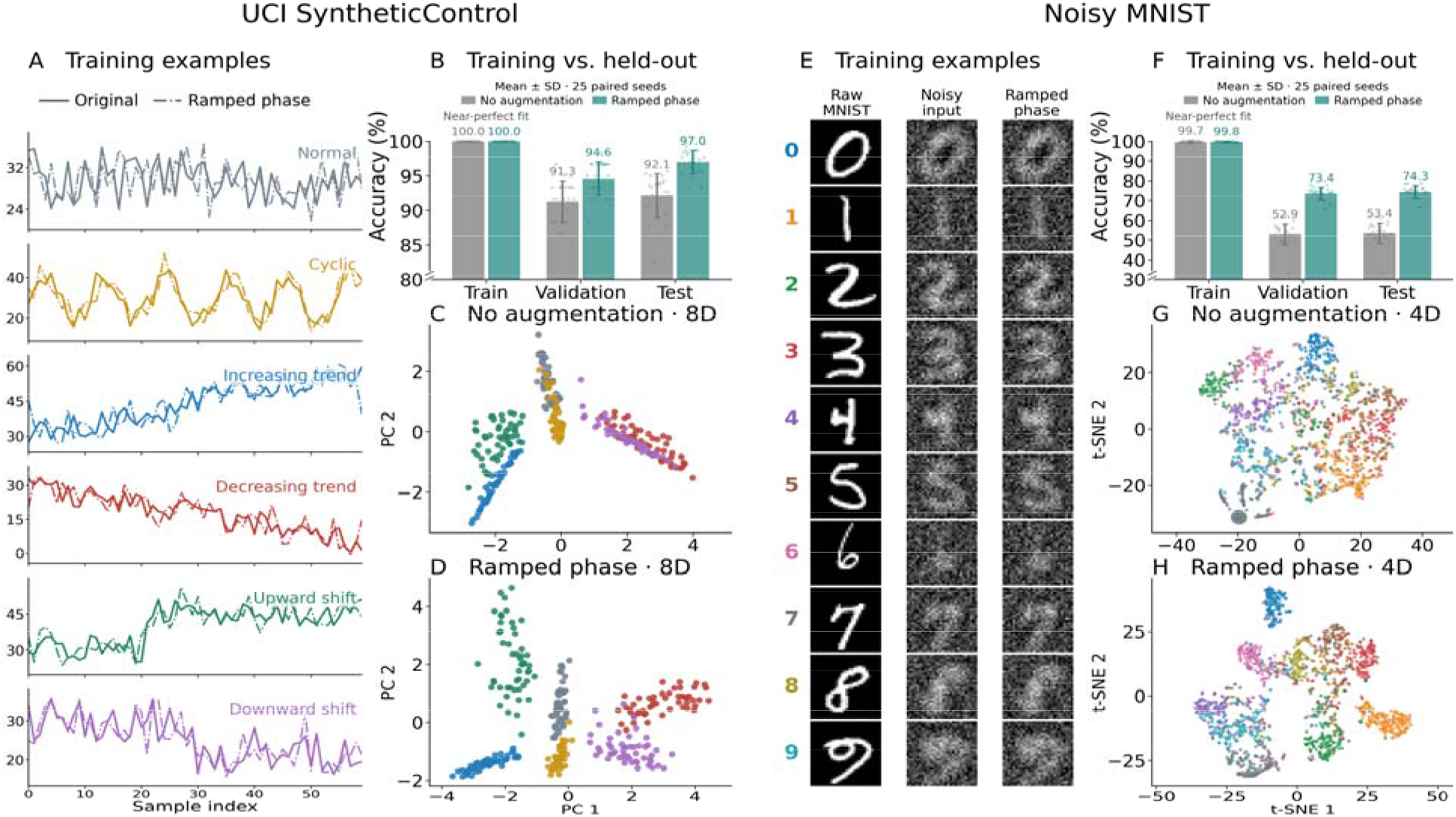
Ramped phase augmentation of two small, noisy training datasets: UCI Synthetic Control (A–D) and noisy MNIST (E–H). (A, E) Example training inputs and one phase-augmented copy of each. (B, F) Training, validation and test accuracy without (grey) and with (teal) phase augmentation for 60 labelled traces and 500 labelled images. Bars show mean ± SD over 25 paired seeds, and points show individual seeds. (C, D) PCA projections of 300 UCI test traces and (G, H) t-SNE projections of the classifier latents of 1,500 MNIST test digits trained without (C, G) and with (D, H) augmentation.

To test the augmentation on images with a known boundary between signal and noise, we constructed a noisy version of MNIST (LeCun *et al*., 1998). Each digit was low-pass limited to 0.16 cycles per pixel, and white Gaussian noise was added at a signal-to-noise ratio of 1. Frequencies above 0.16 cycles per pixel therefore contain only noise (Figure 5E). A classifier with a four-dimensional latent layer was trained on 500 labelled noisy images, 50 per digit, and tested on the 10,000 test digits. Without augmentation, the network classified 99.70% of the training images correctly but only 53.44% of the test digits, a gap consistent with overfitting to the noise of individual images (Figure 5F). Phase augmentation kept training accuracy near 100% (99.81%) and raised mean test accuracy to 74.27% (mean paired gain 20.83 percentage points, 95% interval 18.50–23.16), with higher test accuracy in all 25 seeds. Test digits also formed more distinct clusters in the latent features with phase augmentation (Figure 5G, H). In sweeps over training-set size and latent dimension, phase augmentation increased mean test accuracy and reduced the mean train–test gap in all 36 settings of each benchmark, including settings in which both conditions fitted the training data almost perfectly (Supplementary Figure S2). Supplementary Section S3 summarizes additional comparisons with other augmentations, including the gain from adding phase augmentation to rotation and translation (Supplementary Figure S3), and with controls matched to the phase perturbation.

## 4. Discussion

Ramped phase randomization extends low-pass filtering from amplitudes to phases, so the filtered signal keeps parent detail only to the extent expected at the target resolution. The filter applies to signals of any dimension and was tested here for 1, 2, and 3 dimensions in two broad areas: cryo-EM map comparison and neural network-based machine learning data augmentation for noisy training data.

In cryo-EM, 3D maps at different resolutions are usually compared after amplitude-only low-pass filtering and amplitude scaling (Joseph *et al*., 2020; Warshamanage *et al*., 2022). Ramped phase randomization limits the parent detail that amplitude-only filtering retains beyond the target resolution (Section 3.1; Figure 2) and can be added to existing pipelines without changing registration or scaling. Each filtered map is used as a single realization, because the mean of many realizations converges to a deterministic amplitude filter and loses the phase randomization (Supplementary Figure S1D, E).

In the ribosome example, the FSC shows that the rotated map agrees less with the parent overall, and the difference map shows where, near the subunit interface, as expected for an inter-subunit rotation. The residual of the non-rotated control, not the operator, sets the background against which a conformational change must stand out (Section 3.2). Because the Fourier amplitude matching used here is global and radial, local real-space amplitude scaling (Joseph et al., 2020) might improve contrast matching between the maps. The ribosome pair illustrates the approach rather than establishing general detection limits, and half maps or directional and local envelopes could extend the radial model where resolution varies across a map.

Reference preparation is a related use. Together with independent half-set processing, filtering the initial reference to very low resolution limits model bias from retained phase information (Henderson, 2013). Gold-standard processing alone, however, does not fully protect against bias, for example, from an incorrect initial reference (Sorzano *et al*., 2022). Model bias is a greater risk when using a high-resolution template, as in two-dimensional template matching (Rickgauer *et al*., 2017), where template bias has been documented (Lucas *et al*., 2023; Zhang *et al*., 2025). Limiting the phase information beyond a chosen resolution could reduce that bias. Applying the operator to two-dimensional template matching will be tested in future work.

Phase augmentation turns a few noisy samples into many copies that behave like repeated measurements. Each copy keeps the low-frequency signal of the original, but the high-frequency pattern changes every time, leaving the network no fixed pattern to memorize. Additional UCI experiments refine this view (Supplementary Section S3). Matched white or spectral noise and time warping, which also vary the pattern of each trace, improved accuracy about as much as phase augmentation, as did a deterministic filter with the average effect of the phase offsets, while scaling the overall size of each trace gave little or no gain. The benefit therefore seems to come from weakening the example-specific high-frequency pattern, by varying or attenuating it, rather than from phase randomization as such. Phase augmentation is still the simpler tool: it creates the variation from each sample itself, with the frequency band as its only setting, whereas matched noise must be scaled to the perturbation it imitates.

The benefit held for the CryoBench particles and for both general benchmarks, across training-set sizes and latent dimensions (Figure 4; Supplementary Figure S2). Gains persisted where both conditions fitted the training data almost perfectly, consistent with reduced overfitting rather than better fitting, and more robust generalization. For the CryoBench particles, augmented training beat unaugmented training on twice as many particles in all 16 comparisons. Phase augmentation could therefore potentially reduce the number of labelled particles needed for supervised tasks in cryo-EM, such as classifying particles after consensus refinement and possibly training particle pickers (Bepler *et al*., 2019).

## 5. Methods

### 5.1 Registration and map handling

Each parent map was registered to the lower-resolution map before filtering. The moving map is resampled onto the fixed-map grid using the geometry in the MRC headers. A seven-parameter (3 rotations, 3 positions, and 1 scale) similarity transform is fitted by L-BFGS-B (Byrd *et al*., 1995) to band-limited normalized cross-correlation on copies downsampled to at most 96 voxels per dimension, with a default frequency limit of 1/*R*_*T*_, scale bounds of 0.9–1.1, a 20° rotation radius and translation bounds of 20 voxels (24 for the 70S ribosome). Every registration starts from the best-scoring of 24 principal-axis orientations based on ellipsoid fitting, so the 20° radius bounds only the local refinement around that start, and the fitted transform is applied once at full resolution.

All Figure 2 maps are unmasked. Each pair was registered by band-limited cross-correlation, then resampled onto a common grid and filtered with 50 seeds. Figure 2 shows central sections, with contrast scaled for display only. For Figure 3, all maps were placed on a 128^3^ grid at 2.93 Å/pixel. The two 13.2 Å maps have no deposited half maps, so their reported resolution (FSC = 0.5) was used directly as the filtering target. Both maps are band-limited just beyond that resolution, and amplitude matching carries this limit into the filtered maps.

### 5.2 Phase and amplitude operations

Phase offsets are drawn from a wrapped normal distribution with widths from equations (5) and κ100 (6) and, corrected by equation (7) when a parent resolution is supplied. Offsets are sampled for independent conjugate pairs; self-conjugate coefficients are left unperturbed, and coefficients with *c* < 10^-6^ are set to zero. Each filtered map and augmented example is a single realization; seed averages are used only for comparison with expected values (Supplementary Figure S1). Radial amplitude gains are ratios of shell-mean Fourier magnitudes, capped at 5, with unit DC gain.

### 5.3 FSC and residual statistics

FSCs exclude the DC term and are computed without masks, because a mask shared by both maps mixes Fourier coefficients and can inflate the FSC at high frequency (Van Heel & Schatz, 2005). Input maps are globally standardized to zero mean and unit variance. Difference maps are computed by subtracting the experimental map from each filtered map. The variance ratio compares the rotated and control difference maps within a soft mask covering both low-resolution maps (threshold 0.5). The local root mean square (rms) in Figure 3 uses Gaussian smoothing with a standard deviation of 13.2 Å.

### 5.4 Data augmentation

Augmentation applies equation (2) with unit amplitude response. The offset standard deviation rises smoothly from zero at the start frequency *s*_0_ to σ _max_ at the end frequency *s*_1_,

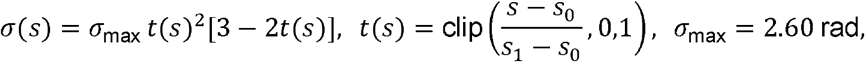

where *s* is a normalized frequency in cycles per sample (time series) or cycles per pixel (images). Each training example is augmented with probability 0.5 at each training step, with new offsets and a start frequency drawn within ±10% of a nominal value; test inputs are not augmented. Reported 95% intervals are t-based confidence intervals for the mean paired difference across seeds, conditional on the fixed test data. Supplementary Section S3 gives the classifiers, seeds and additional comparisons.

For the cryo-EM experiment, we used the deposited Ribosembly particles of the 16 states (128 × 128 pixels at 3 Å per pixel; nominal signal-to-noise ratio 0.01) with the supplied poses, CTF parameters and labels, without adding noise. Training and validation particles came from the first 2,048 particles of each state, with a pool of 1,280 for training and 64 for validation, and the test set was a separate block of 256 particles per state. Each image was shifted by its supplied translation and phase-flipped with the sign of its supplied CTF, with the contrast made positive. The image was then cropped to its central 112 × 112 pixels (336 Å), which contain the whole particle in every orientation, and augmentation was applied to this crop. Each image was backprojected along its viewing direction onto a 144^3^ grid spanning 288 Å (2.0 Å spacing). The selected grid is finer than the 3 Å pixels because, in a tilted view, the corners of the image’s Fourier transform can lie along a grid axis at 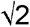 times its Nyquist frequency, and a 3 Å grid would alias them. The volume was multiplied by a soft spherical window (radius 132–156 Å), flattened to 2,985,984 values and standardized with one mean and standard deviation from the training inputs. Training pools, validation and test sets were disjoint within every state (Supplementary Table S1).

Because each first-layer unit sums about three million inputs, one update of the Adam optimizer (Kingma & Ba, 2015) at a learning rate of 0.001 changed the first-layer activations by about 100 times their initial spread, and half of the units stopped responding within five updates. We therefore reduced the learning rate of the first-layer weights to 1.37 × 10□ □, so that one update changes these activations by about a sixth of their initial spread, and kept 0.001 for all other parameters.

### 5.5 Data and software availability

All cryo-EM maps are identified by EMDB accession (The wwPDB Consortium *et al*., 2024) in Figures 2, 3 and S1. The source code for phaselp, which implements registration, filtering, radial amplitude matching, FSC and difference-map analysis with MRC map input and output, is released at https://github.com/jianglab/phaselp. The cryo-EM workflow is also available as a browser-based application at jianglab.github.io/phaselp/. The analysis scripts and figure data are available from the authors upon request. The augmentation experiments use the UCI Synthetic Control Chart dataset (Alcock, 1999) and MNIST (LeCun *et al*., 1998). The cryo-EM augmentation experiment uses the Ribosembly particle images, poses, CTF parameters and maps from CryoBench (Jeon *et al*., 2024), deposited at Zenodo (record 12528292).

## Acknowledgements

This work was supported in part by NIH grants R01NS110437, R01AG071177, R01AG091375, R21GM164827, and Penn State startup funds (WJ). The authors used ChatGPT (OpenAI) and Claude (Anthropic) to assist with writing analysis code and polishing the manuscript text. The authors reviewed all AI-generated content, checked it for errors, and take full responsibility for the published work.

## Supplementary information

### S1. Coherence of wrapped-normal offsets and expected FSC between maps

For offsets drawn from a zero-mean normal distribution with standard deviation σ(s), the coherence of equation (3) is

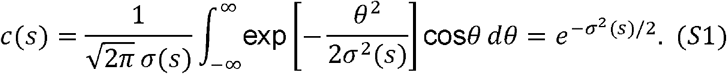

The integral is equivalent to the Fourier transform of a Gaussian, which is again a Gaussian. Wrapping the offsets modulo 2π does not change cos δ. The result also holds for wrapped-normal offsets. Inverting the result gives the third relation in equation (6).

### Expected FSC between independent maps

Let *X* and *Y* be two maps of the same structure *S*, with errors that are uncorrelated with *S* and with each other. In a shell with signal power *P*_*s*_ and error powers *N*_*x*_ and *N*_*Y*_, the cross terms vanish in expectation. For shells with many coefficients, the FSC of each map with the ideal map is therefore 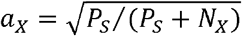, and the FSC between the two maps is

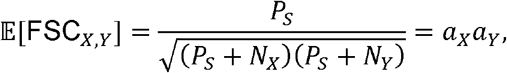

which is equation (8); it holds for any positive scaling of the signal within a shell. For half maps, *N*_*x*_ = *N*_*Y*_, so 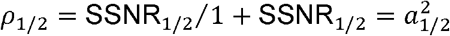. Averaging the two half maps halves the error power, so the full map has twice the SSNR and envelope *a*^2^ = 2ρ _1/2_ / (1 + ρ _1/2_). Equation (5) sets the full-map SSNR to 1/3 at 1/*R*, which corresponds to a half-map SSNR of 1/6 and a half-map FSC of 1/7 ≈ 0.143. For a filtered parent *H*^*’*^, phase offsets independent of the parent multiply its correlation with the ideal map by *c*, so 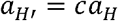, whereas its expected FSC with the parent is *c* (equation 4). With *c* from equation (7), 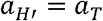, and the expected FSC of *H*^*’*^ with an independent map at the target resolution is 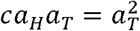.

### Simulation

Figure S1 checks these relations numerically, with the registered T20S parent map (EMD-6287; \(256^3^) grid, 1.288 Å voxels), standardized to zero mean and unit variance, as the ideal map. To link the half-map and full-map criteria, two copies of the ideal map were filtered with equation (2), each with coherence 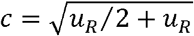 for *R* = 8 Å, where *u*_*R*_ is the full-map SSNR of equation (5), and then averaged (Figure S1A). The copies behave as half maps: their FSC crossed 0.143 at 7.98 Å, each copy correlated with the ideal map at 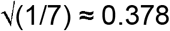 at 8.00 Å, and their average correlated with the ideal map at 0.5 at 8.00 Å (Figure S1B). For maps of different quality (Figure S1C), half maps at resolution were simulated by adding Gaussian noise with shell power to the ideal map, so that their average follows equation (5). A full map at Å, chosen close to the target so that the predictions separate, was filtered to Å with equation (7) and compared with the parent, the ideal map and an independent full map at. Averaged over 20 draws, all simulated FSCs agreed with the predictions to within 0.006 up to 1.5/. Averaging realizations removes the phase randomization (Figure S1D, E). Because the offset factor has mean (equation 6), the mean of independent realizations converges to the deterministic amplitude filter, which keeps the phases of the parent. Its expected FSC with the parent is, which rises from for a single map (equation 7) toward 1. For the ideal map filtered to Å, the means of 1, 4, 16 and 64 realizations agreed with this expression to within 0.009 up to 1.5/ over five draws.

**Figure S1.**
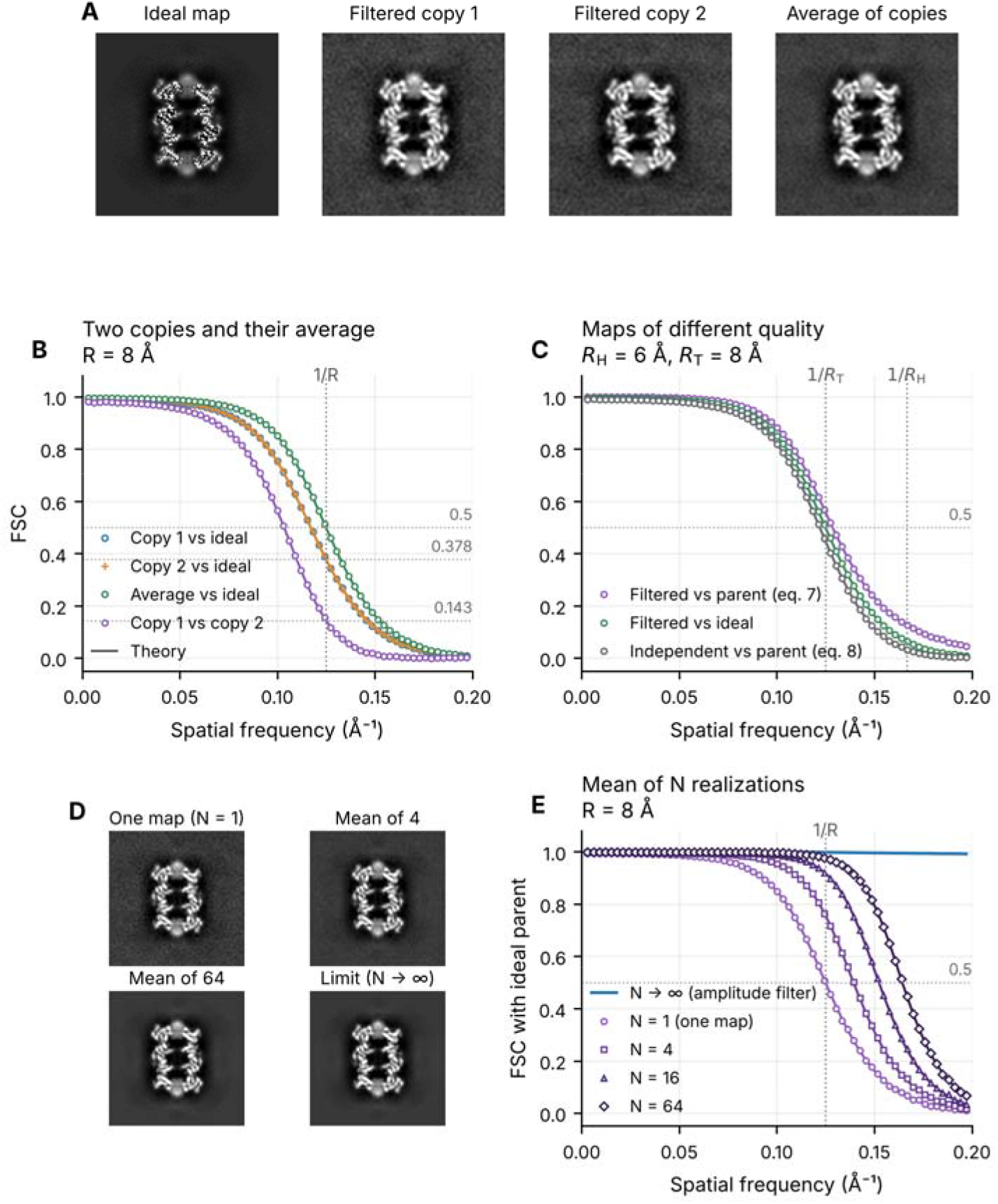
Expected FSCs in simulation. (A) Central sections of the ideal map (T20S parent, EMD-6287), two ramped-phase copies filtered to behave as half maps at Å, and their average. (B) FSC of each copy and of their average with the ideal map, and FSC between the two copies. At 1/*R*, the copies agree with each other at 0.143, while their average agrees with the ideal map at 0.5. (C) Maps of different quality: FSC of a filtered parent (*R*_*H*_ = 6 Å, filtered to *R*_*T*_ = 8 Å) with the parent (*a*_*T*_/*a*_*H*_, equation 7) and with the ideal map (*a*_*T*_), and FSC of an independent full map at *R*_*T*_ with the parent (*a*_*H*_ *a*_*T*_, equation 8). (D) Central sections of a single ramped-phase map (ideal map filtered to *R* = 8 Å), of the means of 4 and 64 independent realizations, and of their limit, the amplitude filter *cT*. (E) FSC of the mean of *N* realizations with the ideal map, which rises from *c* (equation 7) toward 1 as *N* grows. Markers show means over 20 draws (five in E), shading ± SD and lines the predictions. All FSCs are unmasked.

### S2.Expected FSC between a filtered map and an independent map

Phase randomization multiplies each Fourier coefficient of the parent *H* by *e*^*iδ*^ and leaves its magnitude unchanged. The offsets *δ* are drawn independently of any other map *L*, and the expected value of *e*^*iδ*^ is the coherence *c* (equation 6). The expected FSC between the filtered parent *H*^*’*^ and *L* is therefore E[*ρ*_*H*_^*’*^_,*L*_] = *c ρ*_*H, L*_, where *ρ*_*H, L*_ is the FSC of the unfiltered parent with *L*. An amplitude response that is constant within each shell cancels in the FSC, so the prediction also holds after amplitude matching.

### S3.Phase perturbation as a training augmentation

Section 3.3 compares phase augmentation with no augmentation (Figures 4, 5 and S2). This section gives the protocol for that comparison (Table S1) and summarizes additional experiments that compared phase augmentation with alternative augmentations, including geometric augmentation (Figure S3). All experiments use the offset profile of Section 5.4 with unit amplitude response, without registration or amplitude matching, and preserve conjugate symmetry and the DC term. Within each seed, the compared methods share initial weights, data order and augmentation masks. Models are evaluated at the final checkpoint without checkpoint selection, and test inputs are not augmented.

**Supervised experiments for Figures 4 and 5**. Table S1 lists the datasets, models and training settings.

**Table S1.**
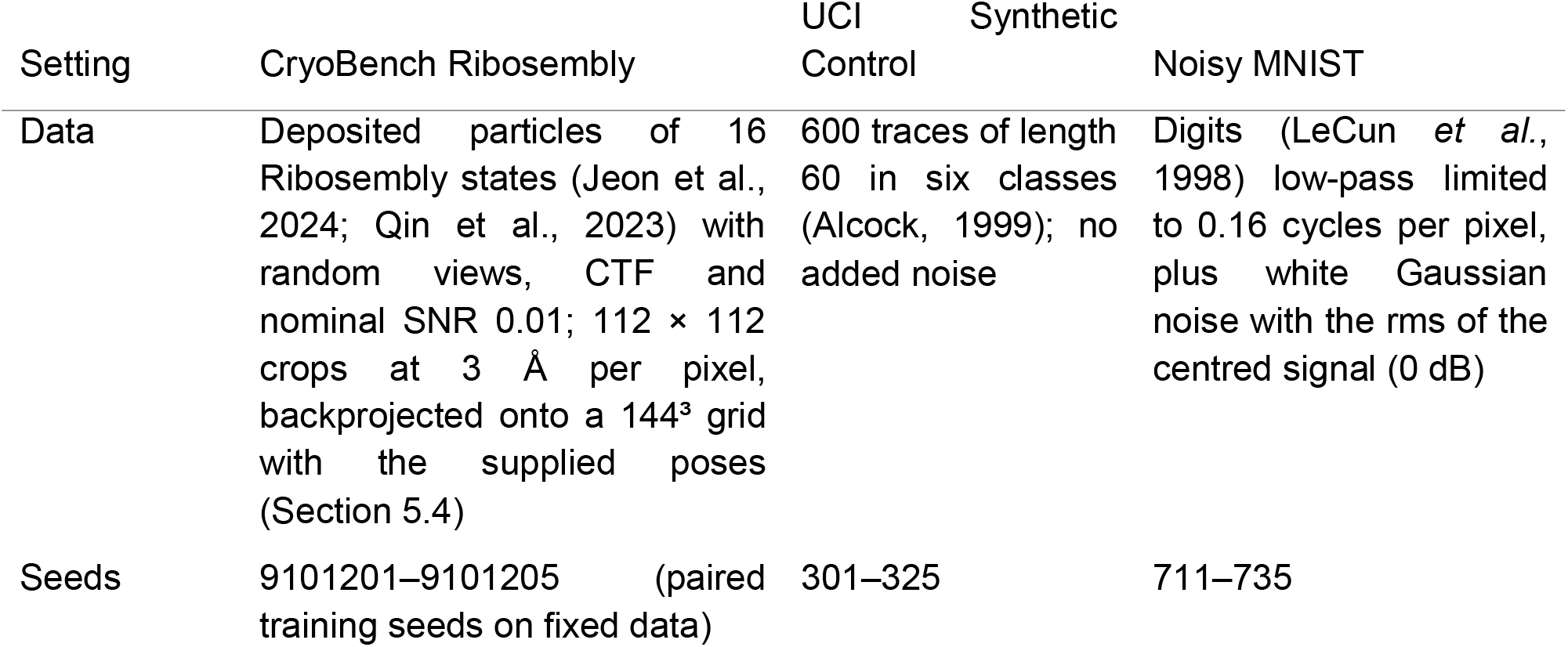
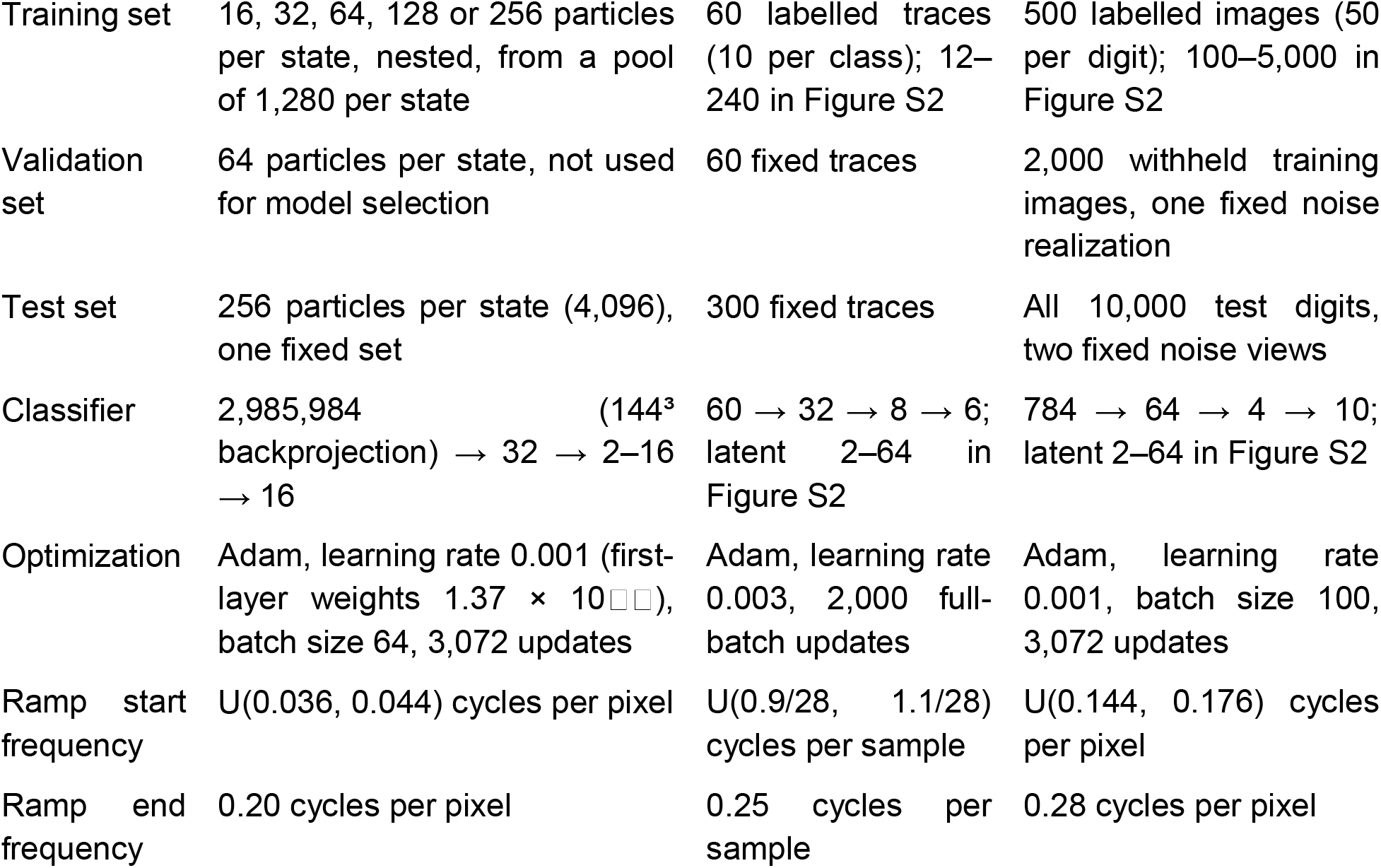
Datasets, models and training settings for the supervised experiments in Figures 4, 5, S2 and S3.

| Setting | CryoBench Ribosembly | UCI Synthetic Control | Noisy MNIST |
|---|---|---|---|
| Data | Deposited particles of 16 Ribosembly states (Jeon et al., 2024; Qin et al., 2023) with random views, CTF and nominal SNR 0.01; 112 × 112 crops at 3 Å per pixel, backprojected onto a 144<sup>3</sup> grid with the supplied poses (Section 5.4) | 600 traces of length 60 in six classes (Alcock, 1999); no added noise | Digits (LeCun *et al.*, 1998) low-pass limited to 0.16 cycles per pixel, plus white Gaussian noise with the rms of the centred signal (0 dB) |
| Seeds | 9101201–9101205 (paired training seeds on fixed data) | 301–325 | 711–735 |
| Training set | 16, 32, 64, 128 or 256 particles per state, nested, from a pool of 1,280 per state | 60 labelled traces (10 per class); 12–240 in Figure S2 | 500 labelled images (50 per digit); 100–5,000 in Figure S2 |
| Validation set | 64 particles per state, not used for model selection | 60 fixed traces | 2,000 withheld training images, one fixed noise realization |
| Test set | 256 particles per state (4,096), one fixed set | 300 fixed traces | All 10,000 test digits, two fixed noise views |
| Classifier | 2,985,984 (144<sup>3</sup> backprojection) → 32 → 2–16 → 16 | 60 → 32 → 8 → 6; latent 2–64 in Figure S2 | 784 → 64 → 4 → 10; latent 2–64 in Figure S2 |
| Optimization | Adam, learning rate 0.001 (first-layer weights 1.37 × 10□□), batch size 64, 3,072 updates | Adam, learning rate 0.003, 2,000 full-batch updates | Adam, learning rate 0.001, batch size 100, 3,072 updates |
| Ramp start frequency | U(0.036, 0.044) cycles per pixel | U(0.9/28, 1.1/28) cycles per sample | U(0.144, 0.176) cycles per pixel |
| Ramp end frequency | 0.20 cycles per pixel | 0.25 cycles per sample | 0.28 cycles per pixel |

### Example and embedding panels

Figure 5 shows seed 303 for UCI, chosen to illustrate the effect, and seed 711 for MNIST. Panels A and E show actual training inputs with one illustrative phase draw each. The UCI PCA is fitted on the training features and applied to the 300 test traces. For MNIST, t-SNE (van der Maaten & Hinton, 2008) embeds 1,500 randomly selected test digits from the first noise view, with perplexity 40 and a fixed initialization seed. Projections use separate coordinates and are illustrations only; the conclusions rest on test accuracy across seeds.

### Cryo-EM particle classification (Figure 4)

Table S1 lists the settings. We trained five paired seeds at every training-set size (16–256 particles per state) and latent dimension (2, 4, 8 and 16), giving 200 models, all tested on the same 4,096 particles. For the ARI, latent features were standardized with training statistics, and k-means with 16 clusters was fitted to the training particles, with each test particle assigned to the nearest centre. The t-SNE panels show seed 9101201 and embed all 4,096 test particles (perplexity 40, 1,500 iterations, fixed initialization seed).

### Training-size and latent-dimension sweeps (Figure S2)

The sweeps used the classifiers, optimizers and ramp settings of Table S1, with latent dimensions of 2, 4, 8, 16, 32 and 64 and new seeds (1301–1325 for UCI and 2301–2325 for MNIST). UCI training sets contained 12, 30, 60, 120, 180 or 240 traces. For each seed, the 600 traces were partitioned anew into 300 test traces, 60 validation traces and a 240-trace training pool, balanced across classes. MNIST training sets contained 100, 250, 500, 1,000, 2,500 or 5,000 images, trained in minibatches of at most 100 images and tested on the 10,000 test digits with the two noise views used for Figure 5. Within each seed, smaller training sets were nested subsets of the largest set, and input normalization used training-subset statistics only.

Phase augmentation increased mean test accuracy and reduced the mean train–test gap in all 36 settings for each dataset, including the 64-dimensional settings not shown in Figure S2.

**Figure S2.**
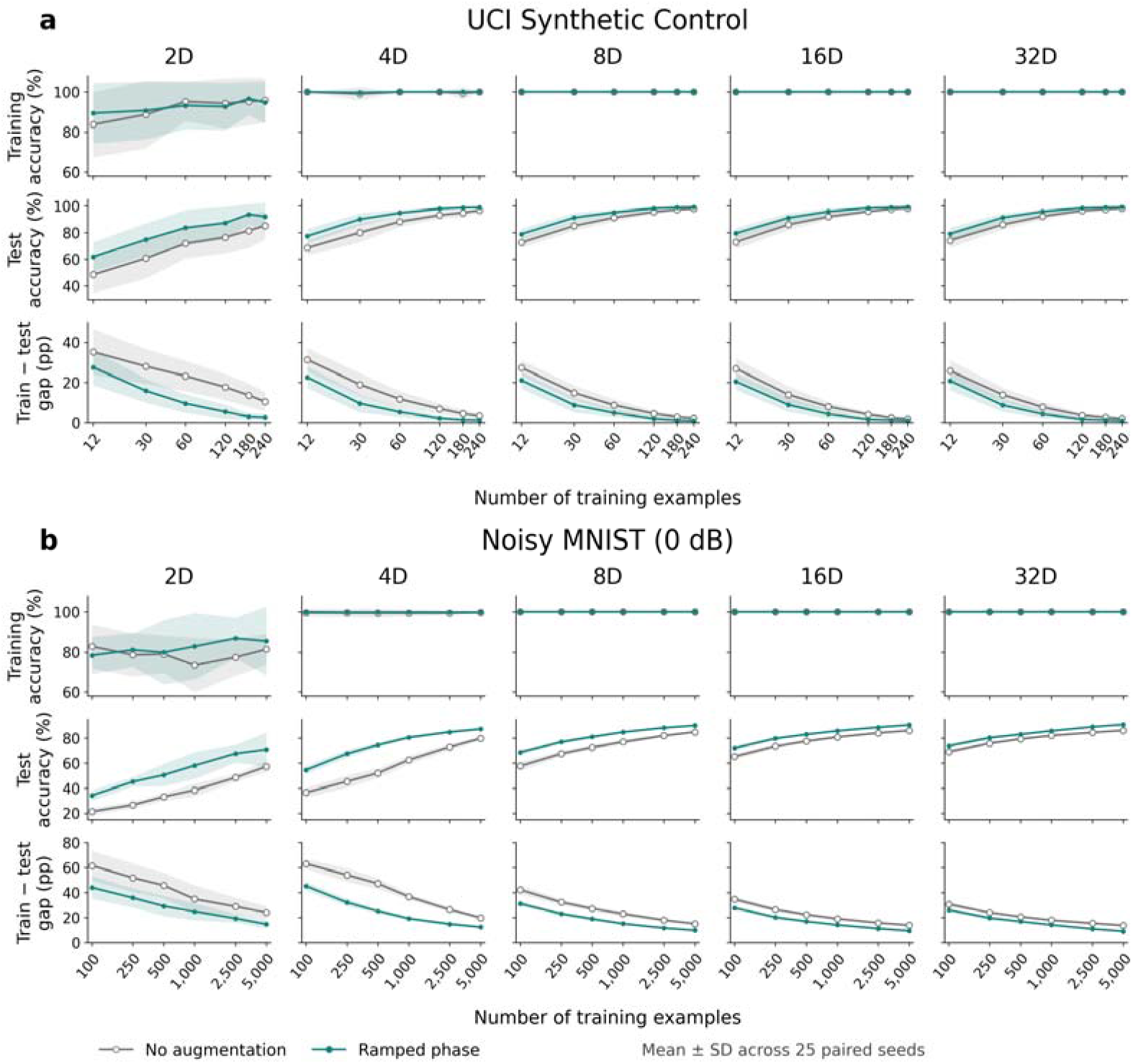
Phase augmentation across training-set sizes and latent dimensions for (a) UCI Synthetic Control and (b) noisy MNIST. Columns show latent dimensions of 2–32, and rows show training accuracy, test accuracy and the train–test gap against the number of training examples on a logarithmic axis. Grey open circles show no augmentation; teal filled circles show ramped phase augmentation. Lines and shading show mean ± 1 SD over 25 paired

Training accuracy, test accuracy and the gap are shown together, because a smaller gap can reflect either better generalization or poorer fitting of the training data. At latent dimensions of 8– 32, both conditions fitted every UCI training set perfectly and every MNIST training set to at least 99.9%. In these settings, mean test accuracy still improved by 0.99–6.43 percentage points for UCI and 3.74–11.03 percentage points for MNIST, consistent with reduced overfitting and more robust generalization. For UCI, gains were largest for the smallest training sets and fell to about one percentage point at 240 traces with 8–32 latent dimensions, where test accuracy without augmentation was already near 98%. For MNIST, the largest gains occurred with two and four latent dimensions, and the setting of Figure 5 (500 images, four dimensions) gave the largest gain in the grid (22.5 percentage points with the sweep seeds). With two latent dimensions, models sometimes underfit, and gap changes there reflect training fit as well as generalization.

### Comparison with geometric augmentation

Geometric transformations have long been used to augment handwritten-digit training data (LeCun *et al*., 1998). To test whether phase augmentation adds to geometric augmentation, the MNIST sweep also used rotation within ±15° and translation within ±2 pixels, alone and after phase augmentation, with matched seeds, training subsets and initializations (Figure S3). The transformations were applied to the noisy training images. Across the 30 displayed settings, phase augmentation alone raised test accuracy by 10.3 percentage points over no augmentation, lower than the 14.3-point gain from rotation and translation. Adding phase augmentation to rotation and translation gave a modest further gain of 0.37 percentage points (averaged over settings; 95% interval across seeds 0.12– 0.62). Mean test accuracy was higher in 24 of 30 settings, including 19 of the 20 settings with 1,000 or fewer training images, indicating a consistent benefit with limited training data.

**Figure S3.**
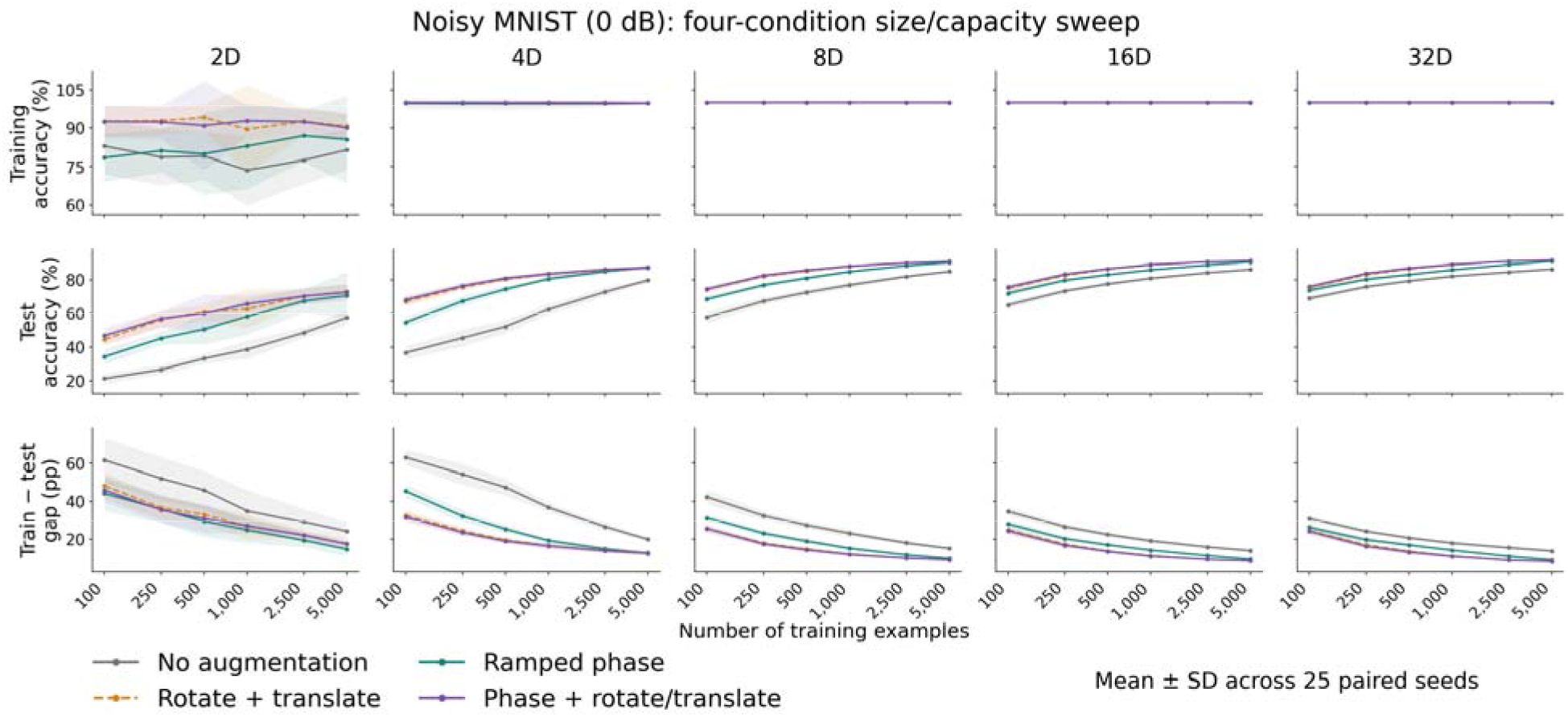
Phase and geometric augmentation in noisy MNIST across training-set sizes and latent dimensions. Columns show latent dimensions of 2–32, and rows show training accuracy, test accuracy and the train–test gap against training-set size on a logarithmic axis. Grey: no augmentation; teal: ramped phase augmentation; orange dashed: rotation and translation; purple: phase augmentation followed by rotation and translation. Lines and shading show mean ± 1 SD over 25 paired seeds. Accuracies are evaluated on unaugmented inputs, and test accuracy averages two noise views within each run.

### Comparisons on UCI Synthetic Control

An additional study used a fixed stratified split (300 test traces, 60 validation traces and a 240-trace training pool), 60 labelled training traces per seed and a 60 →32 →16 →6 classifier trained as in Table S1, with the ramp fixed between 1/28 and 0.25 cycles per sample. Over ten seeds (301–310), phase augmentation was compared with no augmentation, scalar amplitude scaling (factor drawn uniformly from 0.9–1.1), smooth time warping and four controls matched to the phase transform. Matched white noise has the expected total squared perturbation of the phase transform, and matched spectral noise has the same residual power at each frequency. Phase-moment Gaussian multipliers match the conditional mean and covariance of the complex Fourier multiplier, and the mean-phase filter applies the expected multiplier deterministically. Mean test accuracy was 93.07% without augmentation and 97.17% with phase augmentation, a gain of 4.10 percentage points (paired 95% interval 3.16–5.04). Time warping (97.03%) and the four matched controls (96.40–96.90%) gave similar gains, whereas scalar amplitude scaling (93.23%) did not.

### Comparisons on noisy MNIST

An additional unsupervised study trained autoencoders (784 →128 →16 →128 →784) on 4,096 unlabelled noisy images for five seeds (301–305), by pixel mean-squared error with Adam (learning rate 0.001, batch size 256, 3,072 updates). Digits were low-pass tapered from 0.10 to 0.16 cycles per pixel, and Gaussian noise confined to 0.18–0.48 cycles per pixel was added at ten times the centred signal power (−10 dB). The input was an augmented noisy image and the target the unaugmented noisy image, with the ramp fixed between 0.16 and 0.28 cycles per pixel. The 16-dimensional representations of the 10,000 test digits, under two independent noise draws, were standardized, clustered by k-means into ten clusters and scored by the ARI against the digit labels. Phase augmentation raised mean ARI from 0.146 to 0.381 (paired 95% interval for the gain 0.217–0.253), higher than the 0.319 reached with rotation and translation and similar to matched spectral noise (0.379), phase-moment multipliers (0.382) and a known-cutoff low-pass filter followed by PCA (0.370). When the test noise was raised to −20 dB, spectral-noise augmentation gave a higher ARI than phase augmentation (0.281 versus 0.151). Intervals in both additional studies are paired t intervals across seeds, not adjusted for multiple comparisons.

